# Neuronal FOS reports synchronized activity of presynaptic neurons

**DOI:** 10.1101/2023.09.04.556168

**Authors:** Margarita Anisimova, Paul J. Lamothe-Molina, Andreas Franzelin, Aman S. Aberra, Michael B. Hoppa, Christine E. Gee, Thomas G. Oertner

**Affiliations:** Institute for Synaptic Physiology, Center for Molecular Neurobiology Hamburg (ZMNH), University Medical Center Hamburg-Eppendorf, Hamburg, 20251 Germany; Department of Biological Sciences, Dartmouth College, Hanover, NH, 03755 USA; Center for Neuroscience, 1544 Newton Court, Davis, CA 95618; Institute of Pharmacology and Toxicology, University of Zürich, Zürich Switzerland

## Abstract

The immediate early gene FOS is frequently used as a marker for highly active neurons. Implicit in this use is the assumption that there is a correlation between neuronal spiking and FOS expression. Here we use optogenetic stimulation of hippocampal neurons to investigate the relation between spike frequency and FOS expression, and report several surprising observations. First, FOS expression is cell-type specific, spiking CA2 pyramidal neurons rarely express FOS. Second, FOS has a U-shaped dependence on frequency: Spiking at 0.1 Hz is more effective than high frequency spiking (50 Hz) while intermediate frequencies do not induce FOS. Third, the pathway from spiking to FOS is not cell-autonomous. Instead, transmitter release and metabotropic glutamate receptor (mGluR) activation are required and, at 0.1 Hz, FOS is induced independently of CREB/calcineurin/MEK pathways. We propose that FOS does not primarily encode a neuron’s own spike frequency but indicates repeated participation in highly synchronized activity, e.g. sharp wave ripples.

## Introduction

More than 30 years ago, Morgan et al. reported that expression of FOS (Finkel–Biskis–Jinkins murine osteogenic sarcoma virus) protein increased in the brain following seizures (*1*). The ensuing search for activity-dependent genes yielded a large number of upregulated genes. The fastest upregulated (15 – 60 min), known as immediate early genes (IEG), are mostly transcription factors like FOS, which in turn increase expression of proteins involved in neuronal growth and synaptic plasticity (*2*, *3*). In addition to its role as a transcription factor, FOS also has direct effects on lipid synthesis in neurons (*4*). The availability of excellent antibodies popularized the use of FOS as a marker of neuronal activity not only in pathological conditions, but also during learning and memory recall. The creation of a transgenic mouse line using the FOS promoter to express a protein of interest specifically in active neurons was a revolutionary innovation (*5*). Expressing optogenetic tools under FOS control allowed manipulating the activity of a defined subset of neurons. For example, when channelrhodopsin expression is induced in mice during fear conditioning, light-induced reactivation of this subset of neurons produces a strong fear response (*6*, *7*). The functional importance of FOS-positive neurons is not restricted to fear memory; spatial memories are also affected when FOS-positive neurons are inhibited (*8*).

Despite the widespread use of FOS as a marker of active neurons, surprisingly little is known about the relationship between neuronal firing and FOS expression. The *fos* promoter is a point of convergence for several activity-dependent signaling pathways (*9*). Activity-dependent induction occurs when serine 133 phosphorylated calcium/cAMP response element binding protein (pCREB) dimerizes with CREB binding protein (CBP) and binds to the *cre* (cAMP response element) of the *fos* promoter. CREB is phosphorylated by several kinases including cAMP-dependent protein kinase A (PKA), calcium calmodulin dependent kinase IV (CaMKIV), protein kinase C (PKC), and ribosomal S6 kinases (RSKs). A second activity-dependent regulator is the phosphatase calcineurin, which dephosphorylates Rb (Retinoblastoma) and de- represses *fos*. The SRE (serum response element) is a third site that may confer activity-dependence on the initiation of FOS expression. Some of these pathways are primarily activated by calcium, some by growth factors (via receptor tyrosine kinases) or G-protein coupled receptors (GPCRs). In most studies of FOS expression, neurons were activated by chemical stimulation (high K^+^) or by pharmacologically induced seizures. The sodium channel blocker TTX efficiently blocks FOS expression, which led to the conclusion that high-frequency spiking is a key trigger for FOS expression (*10*). Accordingly, long high-frequency bursts of action potentials drive FOS expression in individual CA1 pyramidal neurons (*10*) but are arguably outside the envelope of physiological activity patterns. In cultured neurons, stimuli delivered in repetitive short bursts drive FOS more efficiently than long duration trains of single action potentials, but FOS induction was not correlated with intracellular calcium levels (*11–13*). In vivo calcium imaging studies also found surprisingly weak correlations between the frequency or amplitude of calcium transients and FOS expression in individual neurons during learning (*14–16*). As its complex transcriptional regulation already suggests, FOS seems to be more than a simple reporter of intracellular calcium or spike frequency.

We decided to investigate the relationship between neuronal spiking and FOS expression under highly controlled conditions in rat hippocampal slice culture. We used the red-light-sensitive channelrhodopsin ChrimsonR to drive action potentials in neurons while blocking fast synaptic transmission to prevent spontaneous and propagating activity. As anticipated, high frequency stimulation (50 Hz) drove FOS expression dependent on the number of action potentials, whereas frequencies between 1 and 10 Hz were ineffective. Unexpectedly, we observed the strongest and most consistent FOS expression following stimulation at 0.1 Hz. Pharmacological dissection revealed that FOS induction at all stimulation frequencies required mGluR activation and was therefore not cell-autonomous. Furthermore, glutamate application or chemogenetic activation of Gq was sufficient to induce FOS in the absence of any action potentials (TTX). We conclude that FOS is best triggered by synchronized glutamate release events repeating roughly every 10 seconds. In vivo, such high synchrony events (sharp-wave ripples) occur frequently during periods of quiet rest and slow-wave sleep and have been causally linked to memory consolidation (*17*) and long-term potentiation (*18*, *19*). We propose that FOS and its downstream proteins are specifically upregulated in neurons that repeatedly participate in ripple events, even if they fire only a single action potential per ripple.

## Results

To determine how neuronal activity relates to FOS expression, we needed to precisely induce action potentials (APs) across all hippocampal subfields. To prevent synaptically driven spiking, NBQX, CPPene and picrotoxin were added during optogenetic stimulation to block AMPA, NMDA and GABAA receptors, respectively. Pyramidal cell subtypes differ in their capacitance, input resistance and resting membrane potential (Fig. 1A), leading to a higher firing threshold in CA3 pyramidal neurons (PNs) when driven by somatic current injections (Fig. 1B). For non-invasive spike induction, we virally transduced organotypic slice cultures with the channelrhodopsin ChrimsonR (Fig. 1C). Using 1 ms flashes of a 625 nm LED (Fig. 1D), we found that the light intensity threshold for spike induction was not different for CA1 and CA3 PNs (Fig. 1E). This finding also held true for CA2 neurons and for a blue-light sensitive channelrhodopsin (CheRiff, fig. S1). Apparently, the larger membrane area of CA3 PNs contains proportionally more ChrimsonR channels, compensating for the differences in neuronal size and capacitance (*20*). At a light intensity of 7 mW/mm^2^, we were able to drive AP trains in different hippocampal subfields with high reliability over a wide range of frequencies (Fig. 1F).

**Figure 1:**
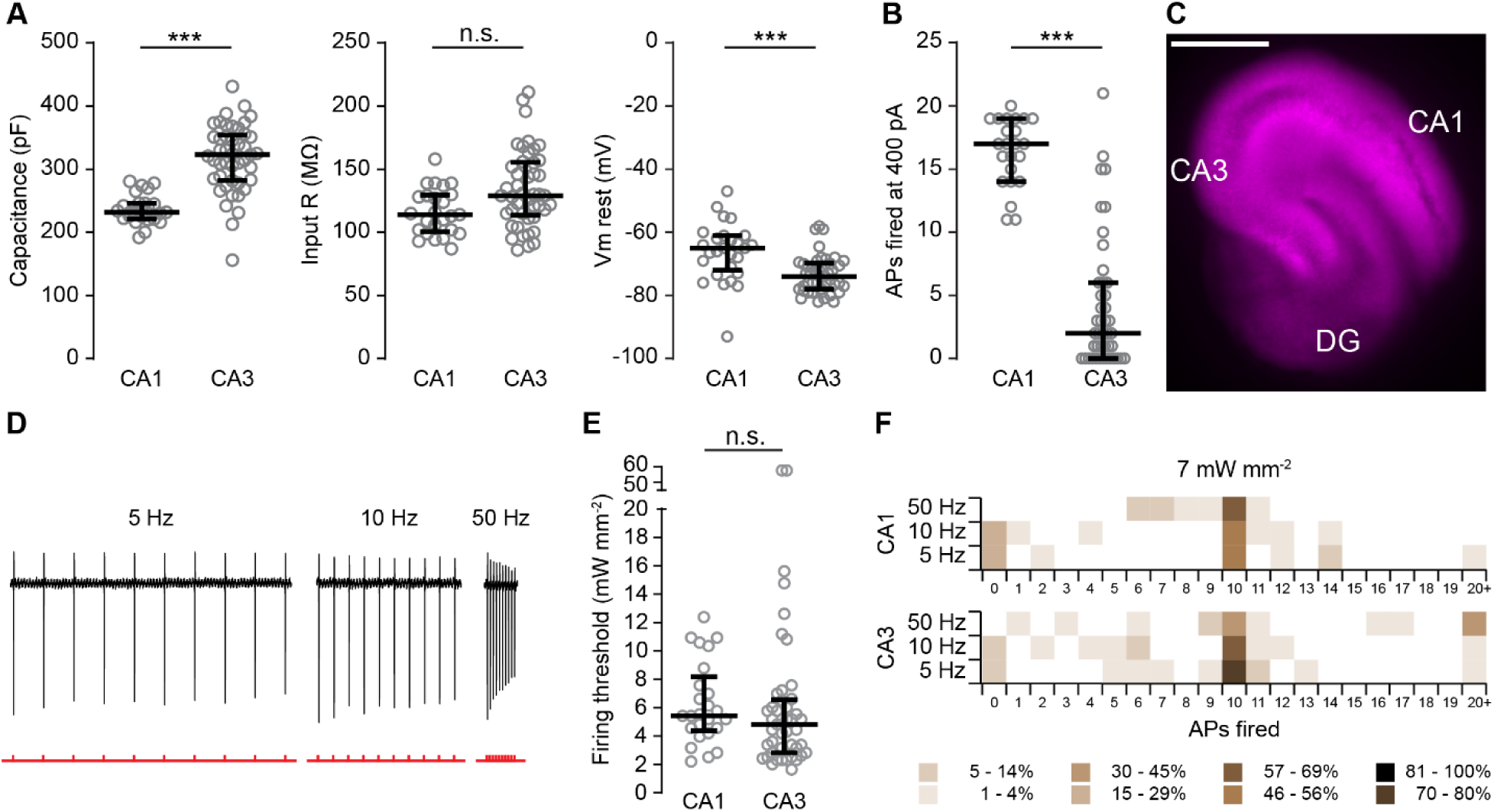
Optogenetic spike threshold is independent of cell size. **(A)** Whole-cell patch clamp recordings from pyramidal neurons in CA1 and CA3 show lower capacitance (*** *P* < 0.0001), similar input resistance (*P* = 0.05) and less negative membrane potential (V_m_; *** *P* < 0.0001) of CA1 pyramidal neurons (CA3: *n* = 45; CA1: *n* = 25; Kolmogov-Smirnov tests). Capacitance and input resistance were measured in voltage clamp, V_m_ in current clamp. **(B)** The number of action potentials (APs) fired during a 600 ms, 400 pA somatic current injection (CA3: n = 44; CA1: n = 25; *** *P* < 0.0001). **(C)** Organotypic hippocampal culture transduced with AAV^2/Rh10^-synapsin-ChrimsonR-tdTomato (magenta). Scale bar 500 μm. **(D)** Optogenetic stimulation of CA1 pyramidal neuron at different frequencies while recording in cell-attached mode. NBQX (10 µM), CPPene (1 µM) and picrotoxin (100 µM) were included to block AMPA, NMDA and GABA_A_ receptors during optogenetic stimulation. **(E)** When driven to spike with light pulses of variable intensity, the firing threshold was not different in CA1 and CA3 pyramidal neurons (CA1: *n* = 25; CA3: *n* = 45; *P* = 0.27, K-S test). **(F)** The majority of pyramidal cells fired exactly 10 action potentials in response to 10 light pulses delivered at 5, 10, or 50 Hz. Plotted are individual data points, median and 25 % to 75 % interquartile range.

As a positive control, we applied high K^+^ (2 x 2 min, 50 mM K^+^ without inhibitors of postsynaptic receptors) and determined that fixing the slices 60 min later is a good compromise for detecting rapidly induced FOS in CA3 and DG neurons as well as the more slowly accumulating FOS in CA1 PNs (fig. S2). By 90 min, FOS immunoreactivity had almost disappeared from CA3 neurons whereas FOS was still increasing in CA1. We next compared the fraction of ChrimsonR^+^ neurons expressing FOS in different regions of the hippocampus after high K^+^ stimulation and after optogenetic induction of 300 APs at 50 Hz inside a cell culture incubator (Fig. 2A). The CA2 region was identified by immunostaining against PCP4/PEP-19 (Fig. 2B), an established marker for CA2 (*21*). Both optogenetic and high K^+^ stimulation produced strong FOS expression in CA1, CA3 and DG, but not in the CA2 region, where significantly fewer neurons expressed FOS (Fig. 2C, D). ChrimsonR expression was homogeneous across all areas and we confirmed that optogenetic stimulation reliably induced APs in all hippocampal subfields (Fig. 1, fig. S1). We conclude that the propensity for FOS expression, and also the time to peak FOS (fig. S2), are systematically different in different cell types. These findings indicate that absence of FOS may not indicate absence of activity, as we have previously observed in DG granule cells (*8*).

**Fig. 2:**
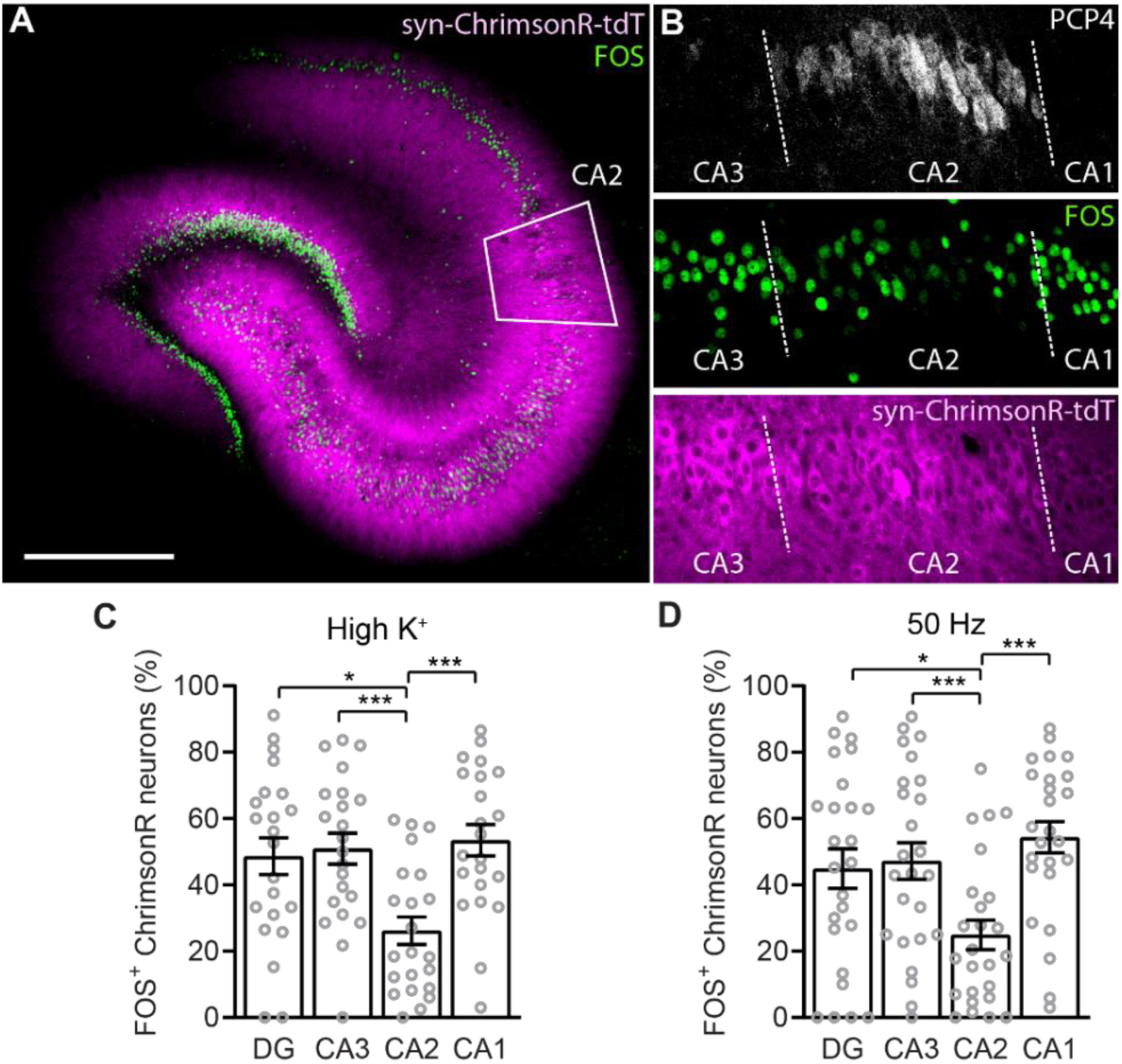
Activity-induced FOS expression depends on cell type. **(A)** A hippocampal slice culture transduced with AAV^2/Rh10^ syn-ChrimsonR-tdTomato (magenta) illustrating widespread high K^+^ stimulated FOS immunostaining (green) in all cell body layers except CA2 (white trapezoid). Slice was fixed 60 min after high K^+^ stimulation. Scale bar 500 μm. **(B)** Single optical section confocal image of a slice stimulated to fire 300 action potentials at 50 Hz, fixed 60 min later and immuno-stained for PCP4 (Purkinje cell protein 4, a marker for CA2 neurons), FOS (green) and ChrimsonR-tdTomato (magenta). White dashed lines indicate the region defined as CA2. Optogenetic stimulation was always performed in the presence of AMPA, NMDA and GABA_A_ receptor inhibitors to block fast synaptic transmission (see Fig. 1). **(C)** The proportion of ChrimsonR-tdTomato-expressing neurons in the dentate gyrus (DG), CA3, CA2 and CA1 that expressed FOS (FOS^+^) one hour after stimulation with high K^+^. Neurons were distinguished from other cell types by tdTomato fluorescence. *n* = 22 slices. Circles correspond to individual slices, bars show mean ± SEM. **(D)** same as C, but stimulation was 300 light flashes, 625 nm, 8 mW mm^-2^ at 50 Hz. *n* = 25 slices. * p < 0.05, *** p < 0.001, RM ANOVA, Tukey’s multiple comparisons.

Next, we explored the number and frequency of APs required for FOS activation. Following in- incubator light stimulation and immunostaining, we counted the number of FOS-expressing neurons in fixed-size ROIs in CA1, CA3 and DG and normalized them to non-stimulated and high K^+^ treated slices to correct for potential variability in the staining of batches (two-point normalization, see fig S3 for non-normalized counts). When stimulated at 50 Hz, FOS expression monotonically increased with the number of light-induced APs (Fig. 3A, B). As 300 light pulses reliably triggered FOS expression, we used this number of pulses to investigate the frequency dependence. To our surprise, the relationship between firing frequency and FOS expression was not monotonically increasing, but strongly bimodal (Fig. 3C). 50 Hz stimulation induced significant FOS expression in DG, CA3 and CA1 (fig. S3), but stimulation at 0.1 Hz was even more effective. To better mimic the stochastic occurrence of synchronized events in the brain, we generated trains of 300 light pulses with Poisson-like statistics (excluding ISIs < 40 ms). Poisson trains with a rate of 0.1 Hz were as effective as regular-spaced 0.1 Hz trains in inducing FOS (Fig. 3D), suggesting that the precise inter-pulse intervals are not critical. As the sodium channel blocker TTX prevented FOS expression at all frequencies (Fig. 3D), induction of FOS required active spiking.

**Figure 3:**
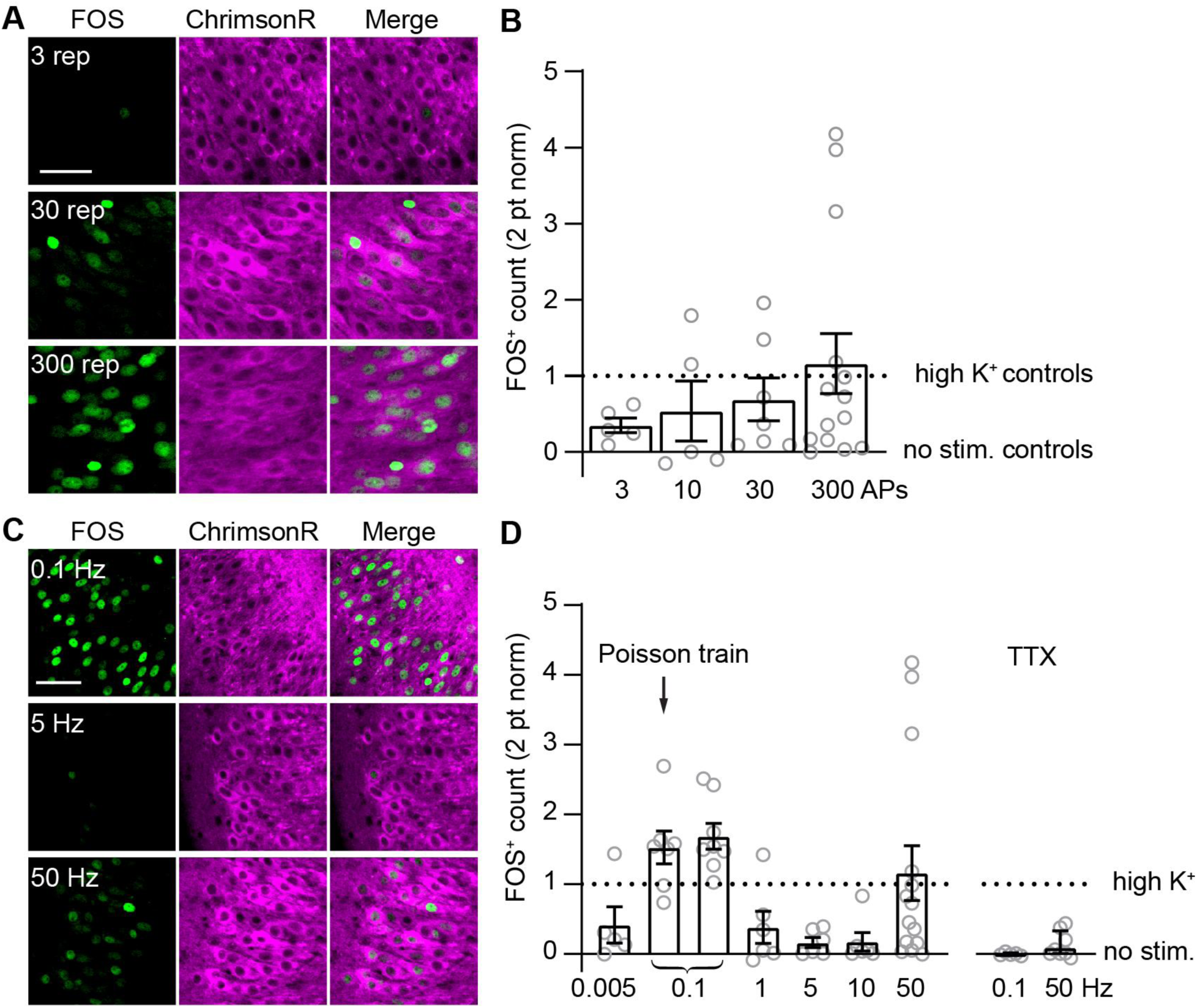
FOS is induced at high and low, not intermediate frequencies. **(A)** CA3 region of hippocampal slices expressing synapsin-ChrimsonR-tdTomato (magenta) and stimulated to fire 3 - 300 action potentials (APs) with AMPA, NMDA and GABA_A_ receptors inhibited. The number of neurons expressing FOS (green) increased with the number of APs. Images are single optical confocal sections. Light stimulation: 1 ms, 625 nm, 50 Hz. **(B)** Total count of FOS^+^ neurons normalized to high K^+^ and non-stimulated control slices included in each batch according to: *x*(*norm*) = (*x* − *min*(*x*))/(*max*(*x*) − *min* (*x*)). Circles correspond to individual slice cultures (*n* = 5, 5, 7, 14), averaging FOS^+^ count in DG, CA3 and CA1. Bars show mean ± SEM. **(C)** As in (A) but stimulated to fire 300 APs at different frequencies. **(D)** As in (B) but experiments as shown in (C) Note that 300 APs at 50 Hz is the same data as shown in (B). At 0.1 Hz the stimulation was applied either regularly or as a Poisson train (3 different trains, minimum interval 40 ms, 2-3 slices each). Tetrodotoxin (TTX 1 µM) blocks voltage gated sodium channels and prevents AP generation. *n* = 5, 7, 8, 6, 6, 6, 14, 5, 8. Scale bars 50 µm.

To better understand the molecular mechanism of FOS induction we targeted three candidate pathways known to induce FOS (Fig. 4A): 1) the phosphoCREB-CBP interaction (666–15), 2) MAPK/ERK - Elk-1/SRF (MEK inhibitor U0126), 3) calcineurin-mediated Rb dephosphorylation (cyclosporin A). As expected, a cocktail of these inhibitors completely blocked FOS induction by high K^+^ and 50 Hz stimulation, while individual inhibitors had less consistent effects against 50 Hz induction (Fig. 4B, C). To our surprise, the same cocktail did not prevent FOS expression after 0.1 Hz stimulation, indicating that other signaling pathways are involved (Fig. 4D).

**Figure 4:**
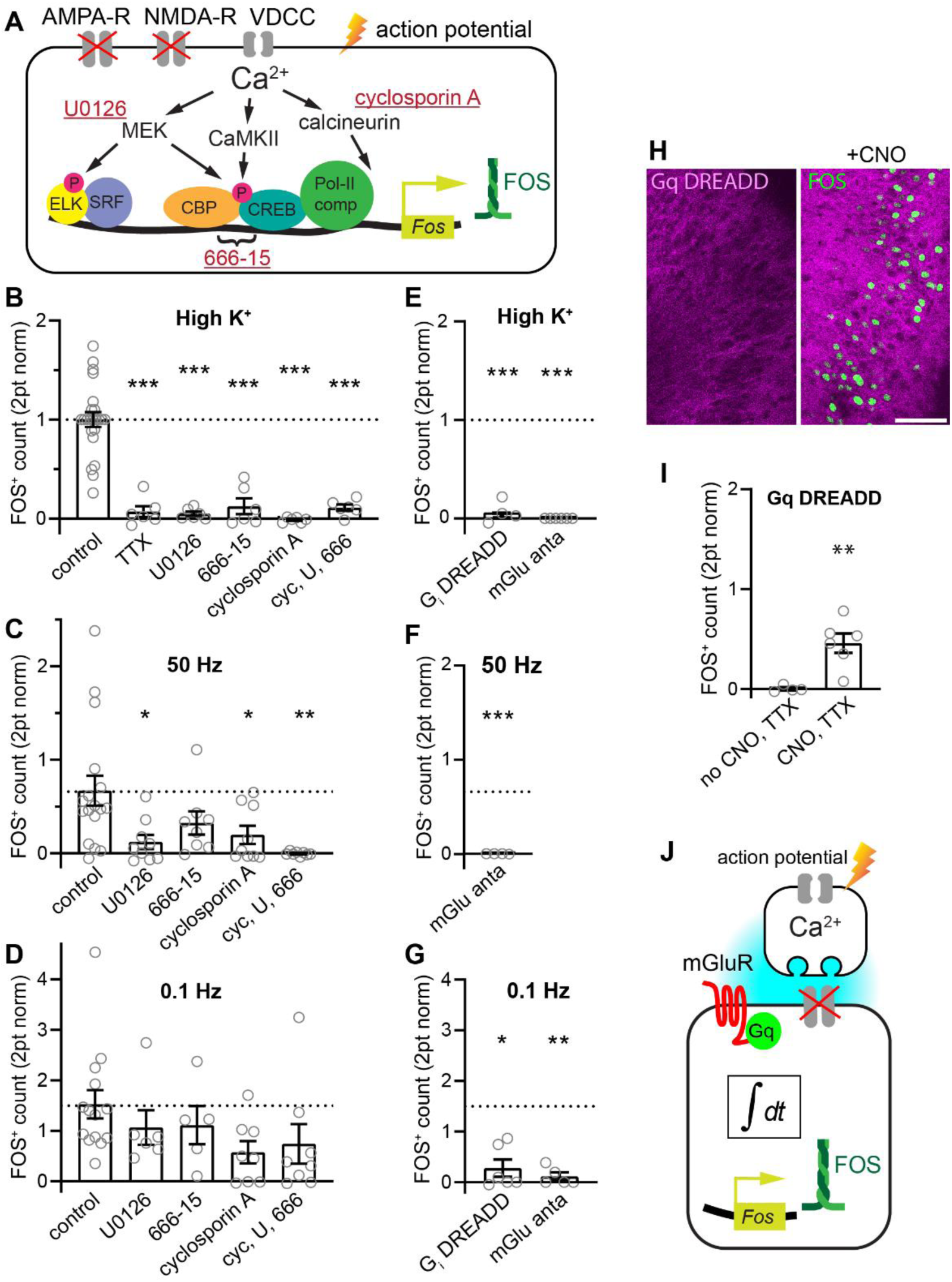
Low and high frequency-induced FOS use different pathways and depend on mGluRs. **(A)** Simplified scheme illustrating the presumed main activity-dependent pathways controlling FOS expression and the inhibitors used. Action potentials trigger calcium influx through voltage-dependent calcium channels (VDCC). U0126 (10 µM) antagonizes mitogen-activated protein kinase kinase ½ (MEK1/2) whose activity leads to activation of serum response factor (SRF) or phosphorylation of cAMP-dependent responsive element binding protein (CREB). 666-15 (666, 800 nM) blocks the interaction of pCREB and CREB binding protein (CBP) which initiates transcription of FOS. Cyclosporin A (cyc 500 nM) inhibits calcineurin which promotes FOS expression by dephosphorylating Rb and releasing transcription (not illustrated). The compounds were applied the evening before stimulating individually or together (cyc, U, 666). **(B)** Number of neurons expressing FOS one hour after stimulation (300 x 0.1 Hz). None of the compounds significantly inhibited FOS induced by 0.1 Hz firing. Normalization to positive (high K^+^) and negative (no stim.) controls as in Fig. 3. Circles correspond to individual slice cultures, bars show mean ± SEM. AMPAR-, NMDAR- and GABA_A_ inhibitors were present. **(C)** Number of neurons expressing FOS after high K^+^ stimulation. FOS expression was inhibited by tetrodotoxin (TTX 1 µM) and by the inhibitors of the pathways illustrated in (A) n = 23, 6, 6, 6, 6, 6. **(D)** As in (B) except slices were fixed one hour after 50 Hz stimulation. n = 17, 9, 8, 9, 8. **(E)** Number of neurons expressing FOS one hour after stimulation to fire 300 APs x 0.1 Hz while transmitter release was inhibited by activation of co-expressed G_i_-DREADD with clozapine-*N*-oxide (CNO) or metabotropic glutamate receptors were antagonized with 2 µM YM298198, 5 µM MPEP and 100 µM MSPG (mGlu anta). AMPAR-, NMDAR- and GABA_A_ inhibitors were present. n = 4, 6 **(F, G)** As E) but after high K^+^ (n = 4, 6) and 300 x 50 Hz stimulation (n = 4). **(H)** CA1 region of slices expressing G_q_DREADD one hour after applying vehicle or CNO. TTX (1 µM) and inhibitors of AMPA-, NMDA- and GABA_A_ receptors were applied the evening before. n = 4, 6. Scale bar 100 µm **I)** Quantification of experiments as in H). **(J)** Key steps in FOS activation missing from the schema in (A). Induction of FOS requires action potentials and glutamate release from presynaptic cells, activation of mGluRs and a temporal integration mechanism (∫*dt*). \**P* < 0.05; \*\**P* < 0.01; \*\*\**P* < 0.001.

Although we blocked activation of fast (ionotropic) postsynaptic receptors, glutamate and other transmitters would still be released by the synchronously spiking neurons. To maintain synchronous spiking but prevent transmitter release, we virally co-expressed ChrimsonR and a Gi-coupled DREADD (*22*, *23*). Under the chemogenetic block of transmitter release, 0.1 Hz stimulation no longer induced FOS, and unexpectedly, even high K^+^-stimulated FOS depended on transmitter release (Fig. 4E, G). This finding strongly argues against the existence of a cell-autonomous spike-driven pathway that leads to FOS expression (*9*, *24*). A parsimonious explanation is that FOS is expressed when postsynaptic neurons receive synchronously released transmitter from many presynaptic neurons. The mGluRs, which respond to both glutamate transients from nearby release sites and spillover from distant synapses, are ideally suitable for detecting such events (*25*). Indeed, neuronal FOS absolutely required mGluR signaling, as antagonizing mGluRs prevented FOS induction by high K^+^, 50 Hz and 0.1 Hz stimulation (Fig. 4E-G).

Group I mGluRs are found near postsynaptic densities and are Gq-coupled. We used a chemogenetic approach to test whether activation of Gq in neurons is sufficient induce FOS in the absence of electrical activity. Activating Gq-DREADD with CNO in the presence of TTX induced FOS in about half of transfected neurons (Fig. 4H, I). Gi activation and all other pharmacological treatments did not induce FOS in non-stimulated cultures (Fig S4). In summary, FOS induction is not a cell-autonomous process (Fig. 4A), but involves communication between neurons via transmitter release (Fig. 4J).

To test the hypothesis that solely firing presynaptic neurons is sufficient to induce FOS in postsynaptic neurons, we made use of the Schaffer collateral pathway. We locally injected ChrimsonR AAV to restrict spiking to CA3 neurons (Fig. 5A to C). With fast synaptic transmission inhibited, stimulation at 0.1 Hz induced FOS throughout the postsynaptic CA1 region (Fig. 5C to G). In contrast, very few ChrimsonR-expressing CA3 neurons expressed FOS (Fig. 5F). This experiment directly demonstrates that FOS is not an indicator of spiking neurons per se, rather, FOS expression indicates which neurons recently received synchronized glutamatergic input.

**Figure 5:**
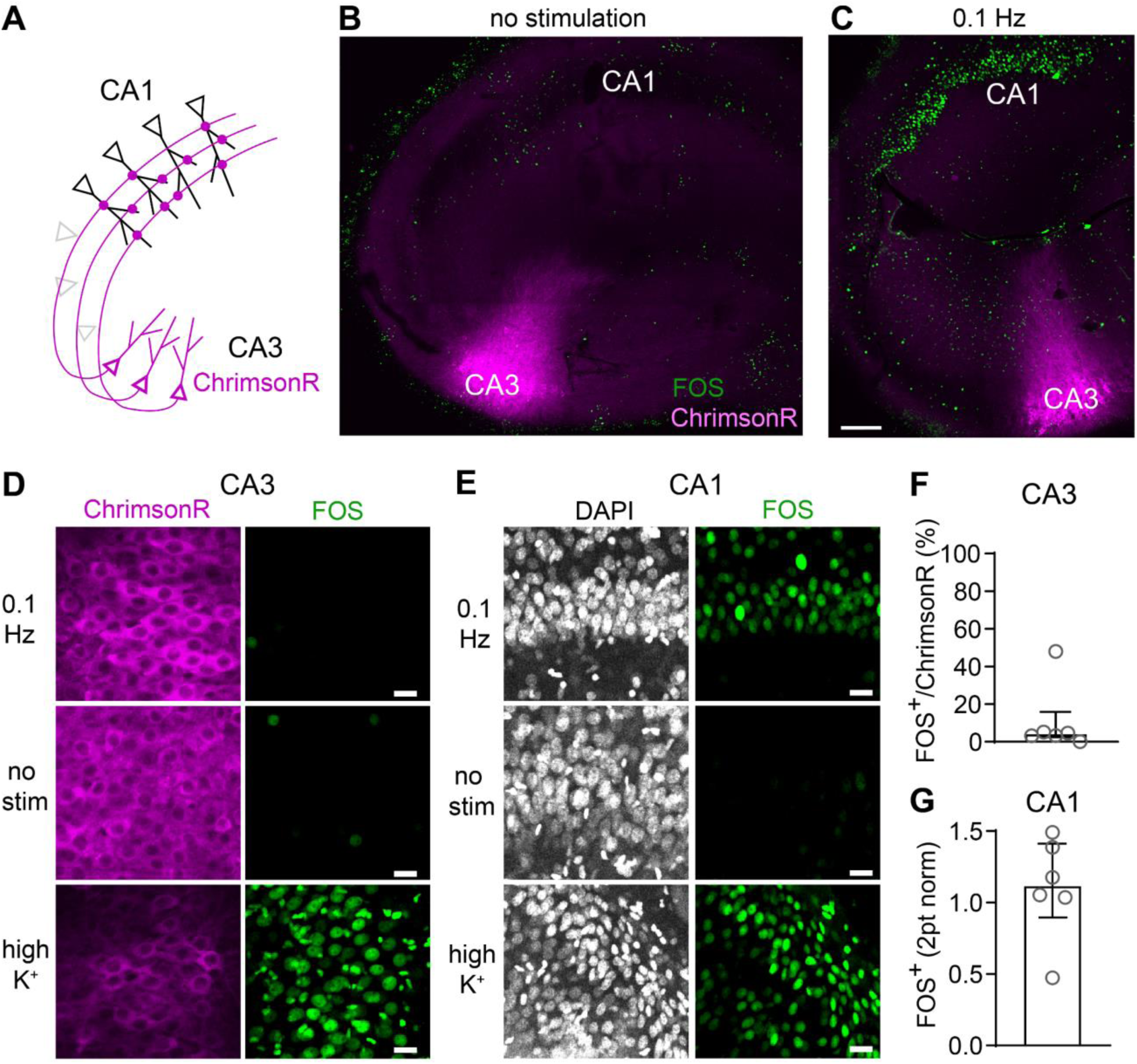
Presynaptic stimulation induces postsynaptic FOS. **(A)** Schema illustrating experiment designed to test whether synaptically released glutamate is sufficient to induce mGluR-dependent postsynaptic FOS. **(B)** Control non-stimulated organotypic culture with a cluster of ChrimsonR-expressing CA3 pyramidal neurons (magenta) stained for FOS (green) expression. **(C)** As in B but stained one hour after stimulation with 300 light pulses at 0.1 Hz. Scale bar: 100 μm applies to B and C. **(D, E)** FOS is not expressed in the optogenetically driven presynaptic neurons (CA3), but in the target region (CA1). Scale bars: 20 μm. **(F)** Following 0.1 Hz stimulation, FOS was expressed in a small fraction of the ChrimsonR- transfected CA3 neurons (3.2%, median with interquartile range, n = 6 cultures). **(G)** FOS expression in CA1 normalized to the non-stimulated and high K^+^ treated cultures (median with interquartile range, n = 6 cultures).

Neuronal postsynaptic mGluRs are localized outside the synaptic cleft, in an annular region (∼100 nm) around the postsynaptic density (*26*). We reasoned that if released glutamate was rapidly diluted from the interstitial space, synchronous spiking would not induce FOS. To create a situation with vastly increased extracellular volume, we applied our induction protocols to dissociated cultured hippocampal neurons in a laminar flow chamber, using electrical instead of optogenetic stimulation (*27*). One hour after the end of stimulation, there was no trace of FOS induction in neurons stimulated at 0.1, 5 or 50 Hz in the presence of antagonists of AMPA and NMDA receptors (Fig. 6A, B). In contrast, directly applying glutamate to stimulate mGluRs strongly induced FOS expression in the cultured neurons (Fig. 6C, D). The dependence on mGluRs was confirmed as there was no effect of inhibiting AMPA, NMDA and GABAA receptors during glutamate application. Similarly, inhibiting fast synaptic transmission did not reduce high K^+^ induced FOS in slice cultures, in line with the requirement for mGluRs (Fig. S5, 4B). As the pan-mGluR agonist L-CCG-1 at both low (10 µM) and high (500 µM) concentrations induced FOS, high affinity (mGlu1-3, 5, 8) rather than low affinity mGluRs (mGlu4 and mGlu7 homomers) are required for FOS expression (Fig. 6C, D).

**Figure 6:**
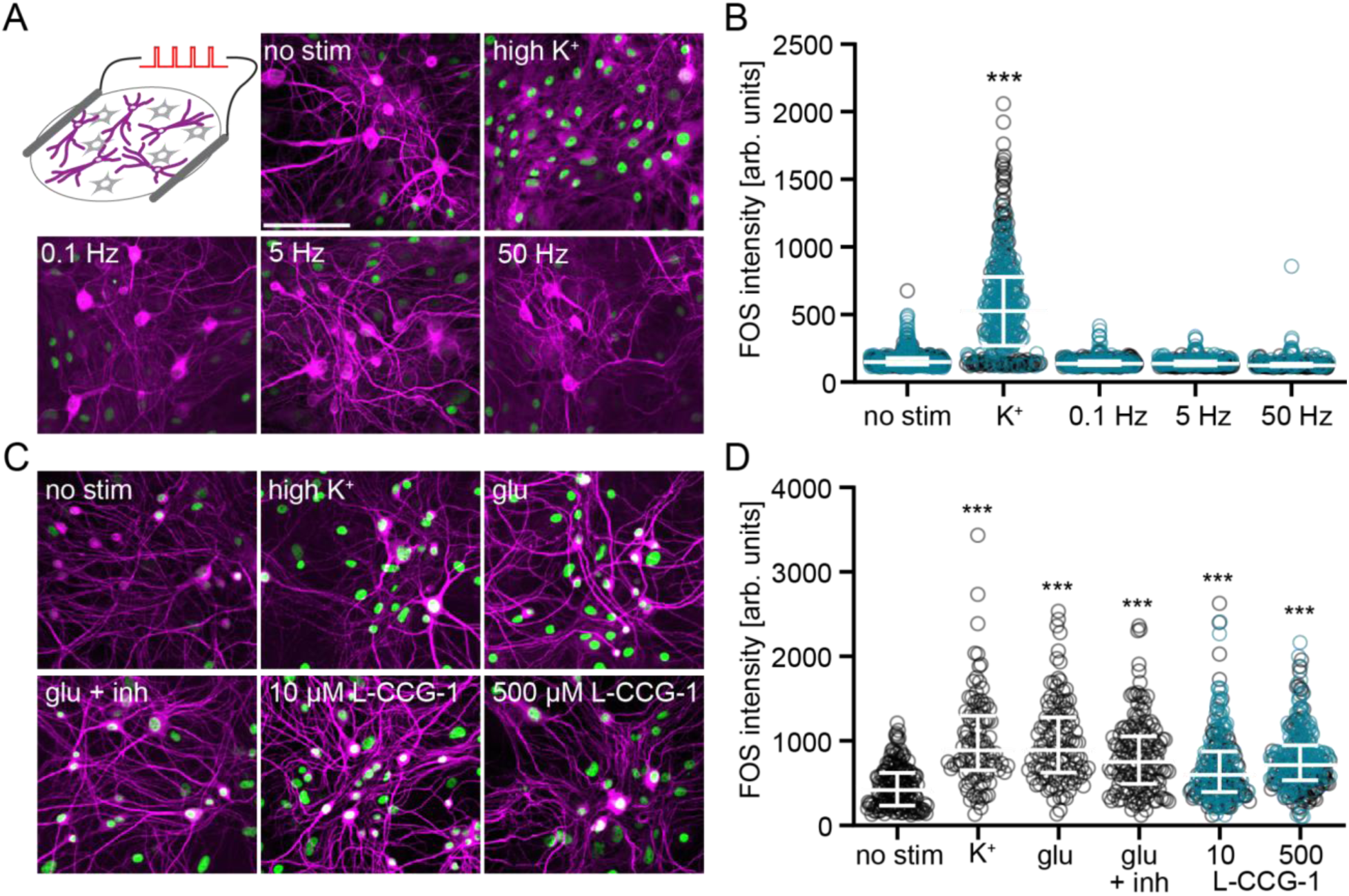
Activation of mGluRs rather than electrically-driven spiking drives FOS in cultured hippocampal neurons. **(A)** Exemplary FOS staining (green) of hippocampal neurons (magenta, MAP2 staining) 1 h after electrical stimulation, high K^+^, or no stimulation (no stim). Note co-localized green and magenta nuclei appear white. **(B)** FOS intensity of MAP2 positive neurons from 2 independent experiments as in A. Circles are intensity of individual neurons, colors indicate different experiments. **(C)** Exemplary FOS staining (green) of MAP2 positive neurons exposed to 50 µM glutamate, 50 µM glutamate + CNQX, APV, picrotoxin to selectively stimulate mGluRs (glu + inh) and the broad-spectrum mGlu agonist L-CCG-1 in the presence of CNQX, APV, picrotoxin to prevent fast synaptic transmission. Different colors indicate independent plates of cultures. **(D)** FOS intensity of MAP2 positive neurons. *** significant increase in FOS compared with no stim (p < 0.0001), Kruskall-Wallis followed by Dunn’s multiple comparisons. Shown are individual values, median and interquartile range. Scale bar 100 µM applies to all panels.

In the high K^+^ experiments, both in slices and dissociated neuronal cultures we noted strong FOS immunoreactivity not only in neurons but also in glia cells (see Figs. 5, 6 and figs. S2, S3). Astrocytes become a powerful source of glutamate under such conditions (*28*) and may contribute to the neuronal mGluR-dependent FOS expression (Fig. 4E).

## Discussion

Our study, which started as a simple calibration attempt to define the relationship between spiking and FOS expression, produced some unexpected results. We expected that a given spiking pattern would similarly drive FOS in all hippocampal principal cells, but we observed that some neurons, i.e., in CA2, express less FOS, whereas the time course of expression differs in others. Surprisingly, the very low frequency of 0.1 Hz stimulation produced more FOS than high frequency (50 Hz) stimulation when the number of action potentials remained at a constant 300. This observation was particularly notable as stimulating the neurons at intermediate frequencies (1 and 5 Hz) almost completely failed to induce FOS. In fact, spiking per se was neither necessary nor sufficient to induce FOS. Rather transmitter release and subsequent activation of mGluRs was necessary, and activation of neuronal Gq signaling was sufficient to induce FOS when spiking was prevented.

Thus, FOS induction is not a cell-autonomous integration of spike-induced calcium influx (*9*, *29*), but a network effect, where individual neurons use GPCRs to detect peaks of extracellular glutamate concentration, integrating this signal over long periods of time (∼1 h). Of course, the same neurons which are receiving and sensing these waves of excitatory glutamatergic input will normally be participating and firing in phase with the input (*30*). The fact that neurons will usually be firing explains how the rather indirect, inter-cellular relationship between activity and FOS has been missed until now. The disconnect between activity (assessed by calcium imaging) and FOS expression at the single cell level has been noted in several studies (*11–13*, *31*). Without the ability to separate postsynaptic from presynaptic spiking afforded by optogenetic tools, however, it was difficult to determine which neurons had to be active to induce FOS (*32*).

As our aim was to define the relationship between simple neuronal spiking and FOS, we purposely inhibited both AMPA and NMDA type glutamate receptors together with GABAA receptors throughout most of this study. Certainly, these receptors contribute to FOS induction either indirectly, by increasing network activity and release of transmitter, or directly by permitting calcium influx (*32*, *33*). Speaking against a direct role of AMPA/NMDA receptors is, however, our observation that FOS induction by high K^+^ was unaffected by their inhibition. Also, in dissociated cell cultures, exogenous glutamate induced the same amount of FOS with and without AMPA/NMDA inhibitors.

In addition to mGluRs, neuromodulators such as dopamine or acetylcholine, via their Gq- coupled receptors, likely contribute to FOS activation during behavior. We speculate these additional receptors may reduce the required fraction of synchronously active presynaptic neurons (and/or the number of repeats) to reach the threshold for FOS induction. Precedents for different types of GPCRs operating in an additive, mutually supporting fashion exist. For example, if mGluRs are blocked, muscarinic M1 receptor agonists restore LTP induced by sharp wave ripple (SWR) type activity (*19*). A candidate mechanism for integrating periodic mGlu activation involves the endoplasmic reticulum (ER). Recently identified mechanisms that link electrical activity with calcium accumulation in the ER (*34*) suggest that the ER could act as a cellular memory of past electrical activity. In this scenario, the magnitude of mGlu-mediated release events would be a function of past activity in that neuron.

What could be the physiological equivalent of our synchronized optogenetic stimulation? Synchrony at 50 Hz (gamma band) does occur during epileptic episodes and indeed leads to massive FOS expression, but is rather unlikely in healthy animals. A prominent activity pattern in the healthy hippocampus are sharp wave ripples (SWRs), high frequency field oscillations inside a short time window (20 – 100 ms) (*17*). Importantly, individual CA1 pyramidal cells typically fire only one or two APs during a single ripple (*35*). SWRs do not occur while the animal is moving, but during quiet wakefulness and slow-wave sleep, more frequently after learning (*17*, *36*). During SWRs, previously active place cells are reactivated (*37*). Strikingly, during periods of immobility or drinking in a maze, mice produce one SWR every 10 seconds, on average (*35*). We propose that our 0.1 Hz optogenetic stimulation may have produced glutamate transients that resemble those experienced by neurons that repeatedly participate in SWRs. Interestingly, oscillations at 0.1 Hz are also prominent in the human electroencephalogram, consistent with cortical synchronization events repeating at this frequency (*38*, *39*).

Replay of activity sequences during SWRs is important for memory consolidation (*18*, *40*). Synaptic potentiation occurs when pre-before-postsynaptic spike sequences are repeatedly combined with ripple-like subthreshold synaptic input (*19*, *41*). Our findings suggest that activation of mGluRs could be the “third factor” promoting LTP induction in these experiments (*42*, *43*). Indeed, it has been shown that FOS^+^ neurons are more strongly connected to other FOS^+^ neurons compared to their neighbors, adding credibility to the idea that FOS may be a marker of the subnetworks that participate in ripples (*44*).

The current study raises some questions which merit further investigation. For instance, are the mGluRs required for FOS induction exclusively located on neurons, or are glial mGluRs also involved in FOS induction (*45*)? How is the activity of metabotropic receptors integrated over long periods of time and what signaling pathway connects the mGluR activation with FOS?

Furthermore, we cannot explain why intermediate stimulation frequencies (1-10 Hz) failed to induce FOS in hippocampal neurons, but are reportedly effective in DRG neurons (*13*). Did these frequencies engage specific FOS suppression mechanisms, e.g. transcriptional repression by DREAM (*46*), or would we eventually see FOS expression if we continue to stimulate at 5 Hz? The parameter space of possible stimulation patterns is almost limitless, but we plan to address some of these questions in future experiments.

Our study has uncovered a GPCR-mediated mechanism that could inform individual neurons whether they are part of a well-synchronized active subnetwork and should ramp up synaptic plasticity (*47*). When using FOS to identify and reactivate “engram ensembles” (*7*, *48*), it is important to keep in mind that these neurons may have never fired at high frequency, but are distinguished by the fact that they participated repeatedly in highly synchronized events. This insight puts a new perspective on optogenetic recall experiments (*6*, *48*), potentially explaining why simple synchronous activation, rather than recreating the precise temporal firing sequence, is sufficient to trigger memory recall.

## Materials and Methods

### Experimental animals

Wistar rats were housed and bred at the University Medical Center Hamburg-Eppendorf (UKE) animal facility and sacrificed according to German Law (Tierschutzgesetz der Bundesrepublik Deutschland, TierSchG) with approval from the Behörde für Justiz und Verbraucherschutz (BJV)-Lebensmittelsicherheit und Veterinärwesen Hamburg and the animal care committee of the UKE. Dissociated neuronal cultures were generated from Sprague–Dawley rat pups (p1) of either sex according to animal protocol number 0002115, approved by Dartmouth College’s Institutional Animal Care and Use Committee.

### Hippocampal slice cultures

The procedure for preparing organotypic cultures (*49*) was modified using media without antibiotics (*50*). Briefly, P5-P6 rats of either sex were anesthetized with 80% CO2 / 20% O2 and decapitated. Using sterile procedures, the hippocampi were removed and 400 µm slices cut on a tissue chopper (McIlwain). Slices were placed on membrane inserts (Millipore, PICMORG50) in 6-well plates and maintained in an incubator at 37°C with 5% CO2. Slice culture medium contained (for 500 ml): 394 ml Minimal Essential Medium (Sigma M7278), 100 ml heat inactivated donor horse serum (H1138 Sigma), 1 mM L-glutamine (Gibco 25030-024), 0.01 mg ml^−1^ insulin (Sigma I6634), 1.45 ml 5 M NaCl (S5150 Sigma), 2 mM MgSO4 (Fluka 63126), 1.44 mM CaCl2 (Fluka 21114), 0.00125% ascorbic acid (Fluka 11140), 13 mM D-glucose (Fluka 49152).

### Primary neuronal culture, stimulation and immunohistochemistry

Primary hippocampal neurons were cultured from hippocampi with dentate gyrus removed, from postnatal day 1 Sprague–Dawley rats of both sexes. Briefly, hippocampal CA1-CA3 regions were digested with trypsin for 5 min at room temperature and dissociated into single cells. Cells were seeded inside a 6-mm-diameter cloning cylinder on polyornithine-coated glass coverslips. For the electrical stimulation experiments, coverslips were mounted in a laminar-flow perfusion and stimulation chamber. Cells were perfused continuously at a rate of 0.4 mL/min in Tyrode’s solution containing the following (in mM): 119 NaCl, 2.5 KCl, 2 CaCl2, 2 MgCl2, 25 Hepes, and 30 glucose with 10 μM CNQX, 50 μM APV, and 100 µM picrotoxin during experiments. APs at 0.1, 5, or 50 Hz were evoked by passing 1 ms current pulses, yielding fields of ∼12 V/cm, through the chamber via platinum/iridium electrodes. All experiments were performed at 34 - 35°C. After 300 APs were evoked, cultures remained in perfusion for an additional hour before fixation. As a positive control, cultures were placed in high potassium (70 mM KCl) Tyrode’s solution lacking synaptic blockers (with equimolar substitution of NaCl) in an incubator for two 5 min periods, after which the cultures were placed back in normal culture media for one hour before fixation. For chemical stimulation, cultures were incubated for two 5 min periods in Tyrode’s containing 50 µM L-glutamic acid (Sigma, G1251) or L-CCG-1 (Tocris, 0333) at 10 or 500 µM. Cultures were then placed in normal culture media with blockers of synaptic transmission (10 µM CNQX, 50 µM APV, and 100 µM picrotoxin) before fixation. Besides the positive controls, these blockers were also added to the cultures 12 h before electrical or chemical stimulation was applied. For immunohistochemistry, primary cultures were fixed in 4% PFA/4% sucrose in PBS for 15 min and washed in PBS (3x). Cultures were then permeabilized with 0.2% Triton X100, washed in PBS (3x), and blocked in 5% goat serum/5% bovine serum albumin (1:1) for 30 minutes. Cultures were incubated overnight at 4°C in primary antibodies (Table 1). The following day, cultures were washed in PBS (3x) and incubated for 1 h at room temperature in secondary antibodies (Table 1) diluted in 5% goat serum in PBS. After a final set of PBS washes (3x), coverslips were mounted for imaging.

**Table 1:** Primary and secondary antibodies.

|  | Primary AB |  |  |  | Secondary AB |  |  |  |  |
| --- | --- | --- | --- | --- | --- | --- | --- | --- | --- |
|  | Anti- | Host | Producer (code) | Dilution | Anti- | Host | Label | Producer (code) | Dilution |
| Fig. 2 | FOS | rabbit | Santa Cruz Inc. (sc-52) | 1:500 | rabbit | goat | Alexa Fluor 647 | Life Tech. (A27040) | 1:2000 |
|  | PCP4 | mouse | Sigma (AMAb91359) | 1:500 | mouse | goat | Alexa Fluor 488 | Invitrogen (A11029) | 1:1000 |
| Fig. 3/4 | FOS | rabbit | Santa Cruz Inc. (sc-52) | 1:500 | rabbit | goat | Alexa Fluor 647 | Life Tech. (A27040) | 1:2000 |
| Fig. 5 | FOS | rat | Syn. Systems (226017) | 1:1000 | rat | goat | Alexa Fluor 647 | Invitrogen (A21247) | 1:1000 |
| Fig. 6 | FOS | rabbit | Syn. System (226-008) | 1:1000 | rabbit | goat | Alexa Fluor 488 | Life Tech. (A11034) | 1:1000 |
|  | MAP2 | guinea pig | Syn. Systems (188-004) | 1:1000 | guinea pig | goat | Alexa Fluor 647 | Life Tech. (A21450) | 1:1000 |
| S2-S5 | FOS | rabbit | Santa Cruz Inc. (sc-52) | 1:500 | rabbit | goat | Alexa Fluor 488 | Live Tech. (A11008) | 1:1000 |

### Viral vectors and transduction

Plasmids containing pAAV-hsyn-ChrimsonR-tdTomato (Addgene # 59171) (*51*) and AAV-hsyn-CheRiff-eGFP (Addgene #51697) (*52*) were packaged into AAV viral particles with serotype Rh10 (7.22×10^13^ vg ml^-1^) at the Vector Core facility of the University Medical Center Hamburg-Eppendorf. For transduction, slice cultures (12-14 DIV) were transferred into sterile pre-warmed (37°C) HEPES-buffered solution containing: NaCl (145 mM, Sigma; S5886-500G), HEPES (10 mM, Sigma; H4034-100G), D-glucose (25 mM, Sigma; G7528-250G), KCl (2.5 mM, Fluka; 60121-1L), MgCl2 (1 mM, Fluka; 63020-1L), CaCl2 (2 mM, Honeywell; 21114-1L), pH 7.4, 318 mOsm kg^-1^. A glass pipette was loaded with virus-containing buffer. For global transduction, a drop of virus-containing buffer from the tip of glass pipette was placed on top of the culture, and for local transduction, glass pipette was inserted into CA3 region under visual control for pressure injection (PicoSpritzer III).

### Optogenetic stimulation in the CO2 incubator

A closed 35 mm petri dish containing the membrane insert with a single slice culture in the center was placed on top of a 625 nm LED (Cree XP-E red) with collimation lens (Roithner LaserTechnik, 10034) as described (*53*). The LED was controlled from outside the incubator by a Grass S8800 stimulator, a constant-current driver (RECOM RCD-24-1.20) and a timer. LED power was adjusted to provide 7 mW mm^-2^ in the specimen plane. For Poisson stimulation at a rate of 0.1 Hz, we generated pulse trains with random inter-pulse intervals (ISIs) in Matlab, excluding ISIs < 40 ms to limit the peak frequency to 25 Hz. TTL pulse trains were sent to the LED driver via a DAQ board (PCIe-6321, National Instruments).

### Slice culture immunohistochemistry

Slice cultures were fixed (30 min in ice-cold 4% paraformaldehyde in PBS) 60 min after stimulation. Slice cultures were washed in PBS (3 x 10 min) and blocked (2 h, 5% donor horse serum, 0.3 % TritonTM X-100 in PBS at RT), then incubated overnight with the primary antibodies (Table 1) in carrier solution (1% Bovine serum albumin, 0.3% TritonTM X-100 in PBS) at 4°C. On the next day, slice cultures were washed 3 x in PBS and incubated at room temperature for 2 h with secondary antibodies (Table 1) in carrier solution (same as above), then washed again (3 x 10 min) in PBS and mounted with Shandon Immu-Mount (Thermo Scientific; 9990402). Figure 5: slices were additionally stained with 4′,6-diamidino-2- phenylindole (DAPI, 1:1000) for 5 min during the last washing step.

### Confocal microscopy

The mounted slice cultures were imaged on an Olympus F1000 confocal microscope using a 20x oil immersion objective and 405, 488, 559, or 635 nm excitation. The imaging parameters were adjusted for each set of experiments and kept unchanged throughout. Z-series (5 slices) stacks with 3 µm z-step at a 1024 × 1024 pixel resolution (zoom 1x) scanning at 12.5 µs per pixel were taken and analyzed in IMARIS software. Fiji/ImageJ was used to project the z-series and overlay channels. The mounted dissociated cultures were imaged on a Nikon Eclipse Ti spinning disk confocal microscope at 2048 × 2048 pixel resolution (20x air objective). Z-stacks with 0.9 µm steps were taken at 3-4 locations within the culture. Fig. 5B, C: Multiple stacks with 50 - 60 µm total depth (3 µm z-step) were imaged (512 × 512 pixel resolution, zoom 1x) with a scanning speed of 4 µs per pixel. The acquired stacks were stitched using Fiji/ImageJ. Fig. 5D, E: Single plane images (1024 × 1024 pixel) were acquired, scanning at 100 µs per pixel.

### Image analysis

IMARIS software was used to count FOS^+^ nuclei and to measure their intensity. FOS^+^ spots (8 µm for CA1 and DG, 10 µm for CA2 and CA3) were automatically detected using the same quality filter threshold throughout all experiments. FOS^+^ spots detected outside the pyramidal cell layer and FOS^+^ nuclei of glial cells were manually removed. For ChrimsonR^+^ neurons with no FOS signal, spots were manually added. Fig. 5: CA1: Six circular regions of interest (ROI, radius 30 µm, z-depth 9 µm) were placed on the CA1 pyramidal cell layer based on the DAPI signal. Within each ROI, all DAPI^+^ neurons were manually selected (8 µm spots). In CA3, ChrimsonR^+^ neurons were manually selected (8 µm spots). Fig. 6: All MAP2^+^ cells were manually selected (8 µm spots). During manual spot addition, the FOS channel was turned off to avoid bias. A spot was defined as FOS^+^ if the intensity was above a specific threshold defined by the negative control (no stimulation) images for each batch of experiments. The absolute number of FOS^+^ neurons in each slice culture was two-point normalized to the number of FOS^+^ neurons in positive (high K^+^) and negative (no stimulation) controls.

## Acknowledgments

We thank Iris Ohmert, Jan Schröder for excellent technical support, Ingke Braren of the UKE Vector facility for preparing the AAVs and Jack Mellor for helpful comments on the manuscript. pAAV-Syn-ChrimsonR-tdT was a gift from Edward S. Boyden, AAV-hsyn-CheRiff-eGFP was a gift from Adam E. Cohen.

## Author contributions

Conceptualization: T.G.O. and C.E.G. Investigation and visualization: M.A., P.J.L.-M., A.F., and A.S.A. Funding acquisition and supervision: T.G.O., C.E.G. and M.B.H. Writing—original draft: T.G.O. and C.E.G.; review and editing: M.A., P.J.L.-M., A.F. and M.B.H.

## Data and materials availability

All data needed to evaluate the conclusions in the paper are present in the paper and/or the Supplementary Materials.

## Supplementary Materials

Figs. S1 to S5.

## Supplementary Material

**Figure S1:**
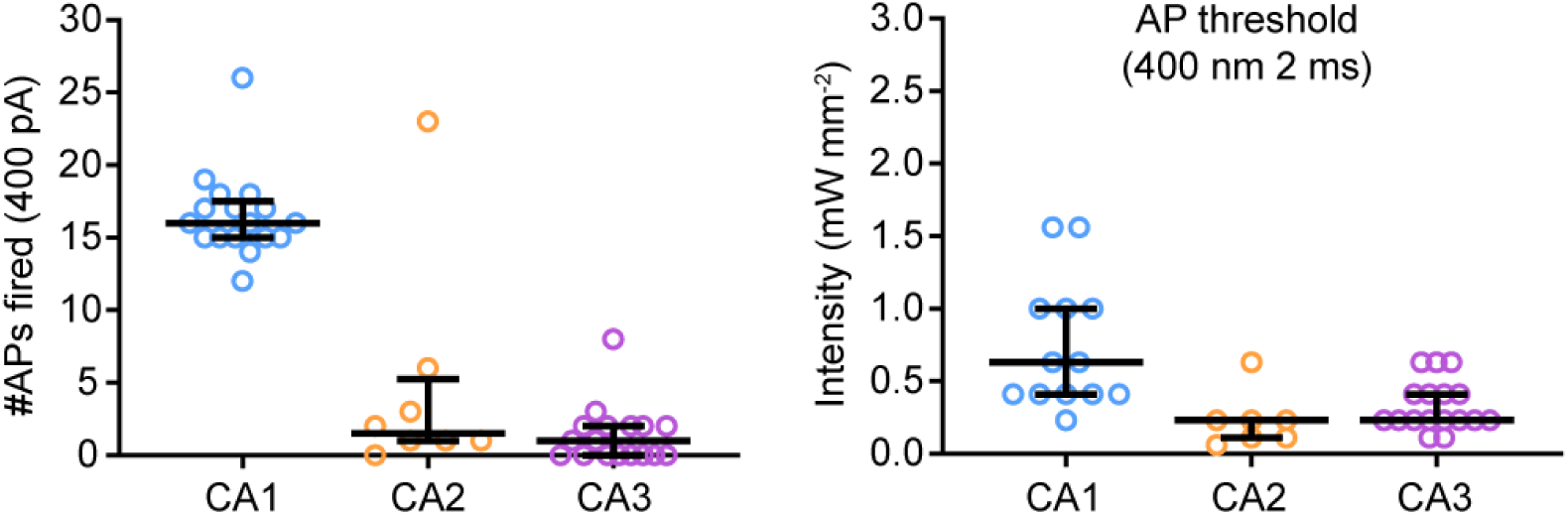
Action potential generation by current injection and light intensity threshold for action potential generation in CheRiff expressing hippocampal neurons. As in figure 1 except hippocampal slice cultures were transduced with AAV2/9-synapsin-CheRiff-eGFP. Light intensity threshold for action potential (AP) was determined in cell-attached mode. The number of action potentials fired in response to a 600 ms, 400 pA current injection from -70 mV was obtained after achieving whole-cell access. n = 17, 8, 19 neurons, respectively. Bars show median ± quartiles. Note that for some cells whole cell-access was obtained before light was applied, so the AP threshold was not measured.

**Figure S2:**
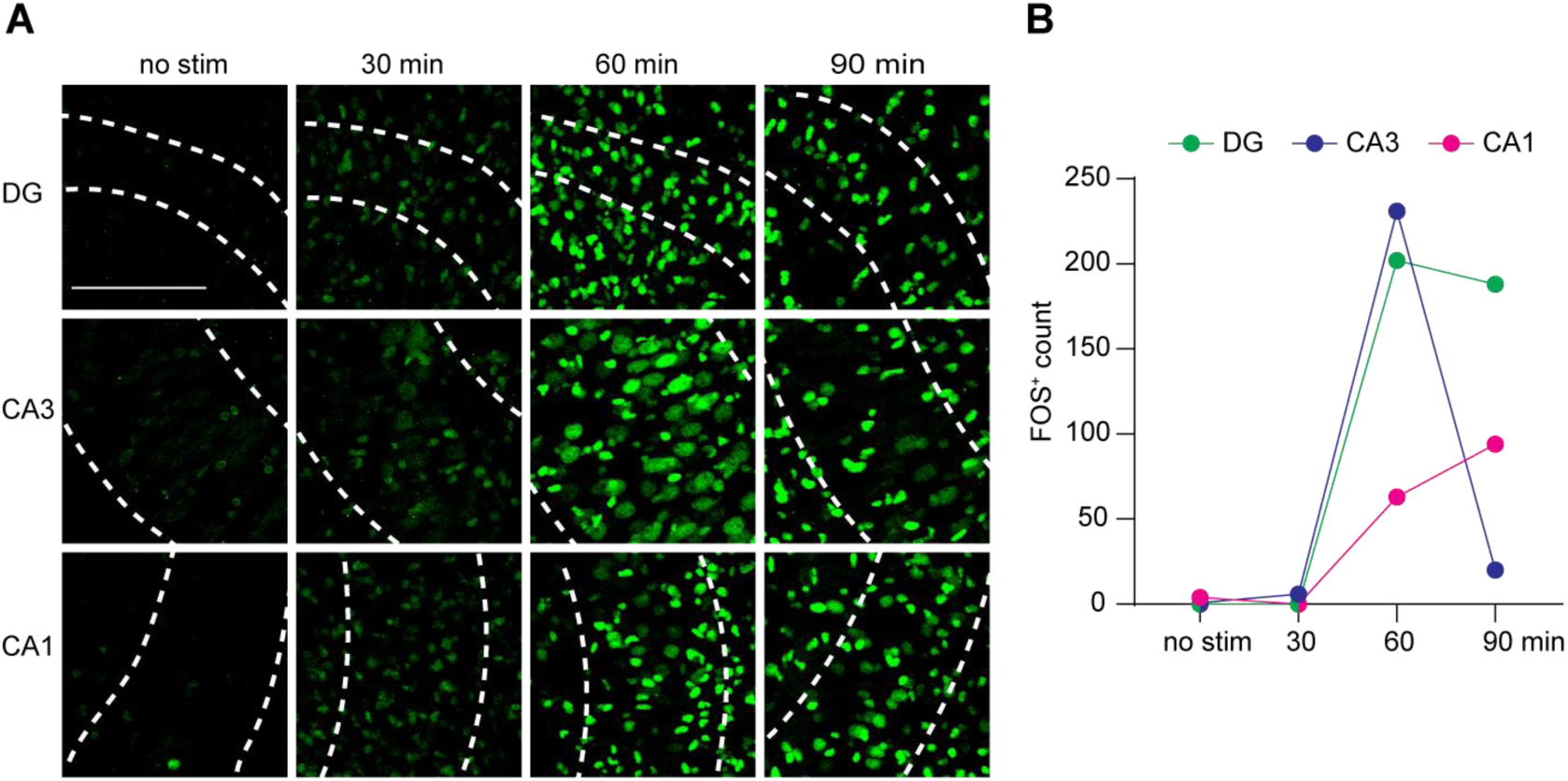
Time course of FOS expression after high K^+^. **A)** Organotypic slice cultures of rat hippocampus fixed at different time points after K^+^ stimulation and stained for FOS (green). Images are maximum intensity projections of 5 optical sections. Scale bar 100 µm. no stim = slice fixed without high K^+^ stimulation. **B)** Count of FOS^+^ neuronal nuclei in hippocampal subfields from slices fixed at different time points. 60 min was chosen as the optimal time point to fix slices after stimulation as expression peaked at this time point in dentate gyrus and CA3 and at 90 min FOS had almost disappeared from CA3.

**Figure S3:**
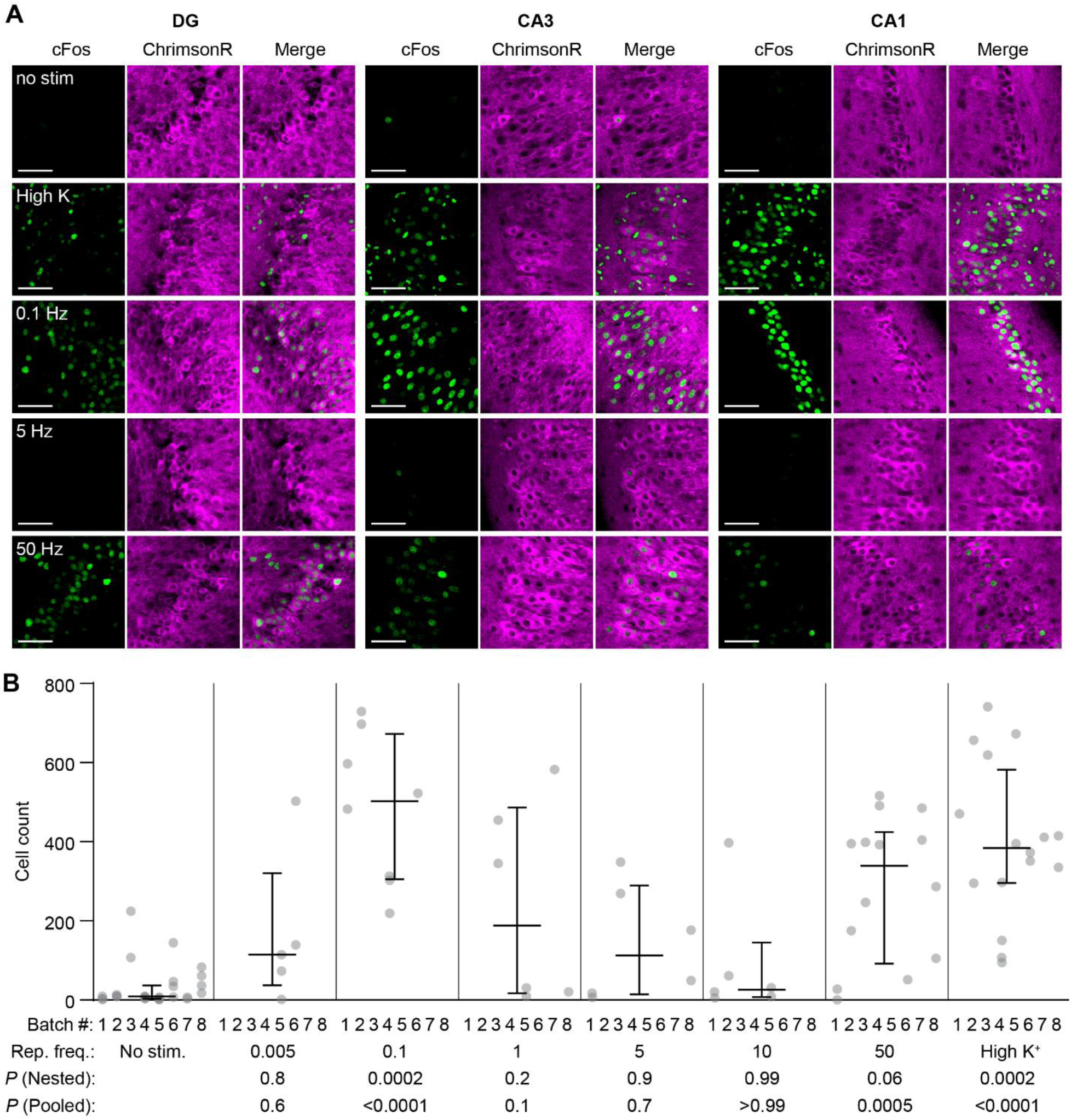
Bimodal relationship between firing frequency and FOS expression. **A)** Confocal images (single plane) of FOS expression pattern 60 min after light stimulation (300 pulses) at different frequencies in 3 hippocampal regions. Magenta: ChrimsonR-tdTomato. Green: anti-FOS. Scale bars: 50 µm. **B)** Quantification of FOS-positive neurons (88 cultures, 8 stimulation conditions, n = 8 biological repeats). Each dot represents one organotypic culture, averaging 3 FOVs (635 x 635 μm) in DG, CA3 and CA1, respectively. In addition to individual data points, median and 25 % to 75 % interquartile range are indicated. *P* (Nested): One-way ANOVA on nested data. *P* (Pooled): Kruskal-Wallis test on pooled data. *P* values are in comparison to ‘No stim.’ group. This set of experiments (stimulation + staining + imaging) were performed in 8 batches.

**Figure S4:**
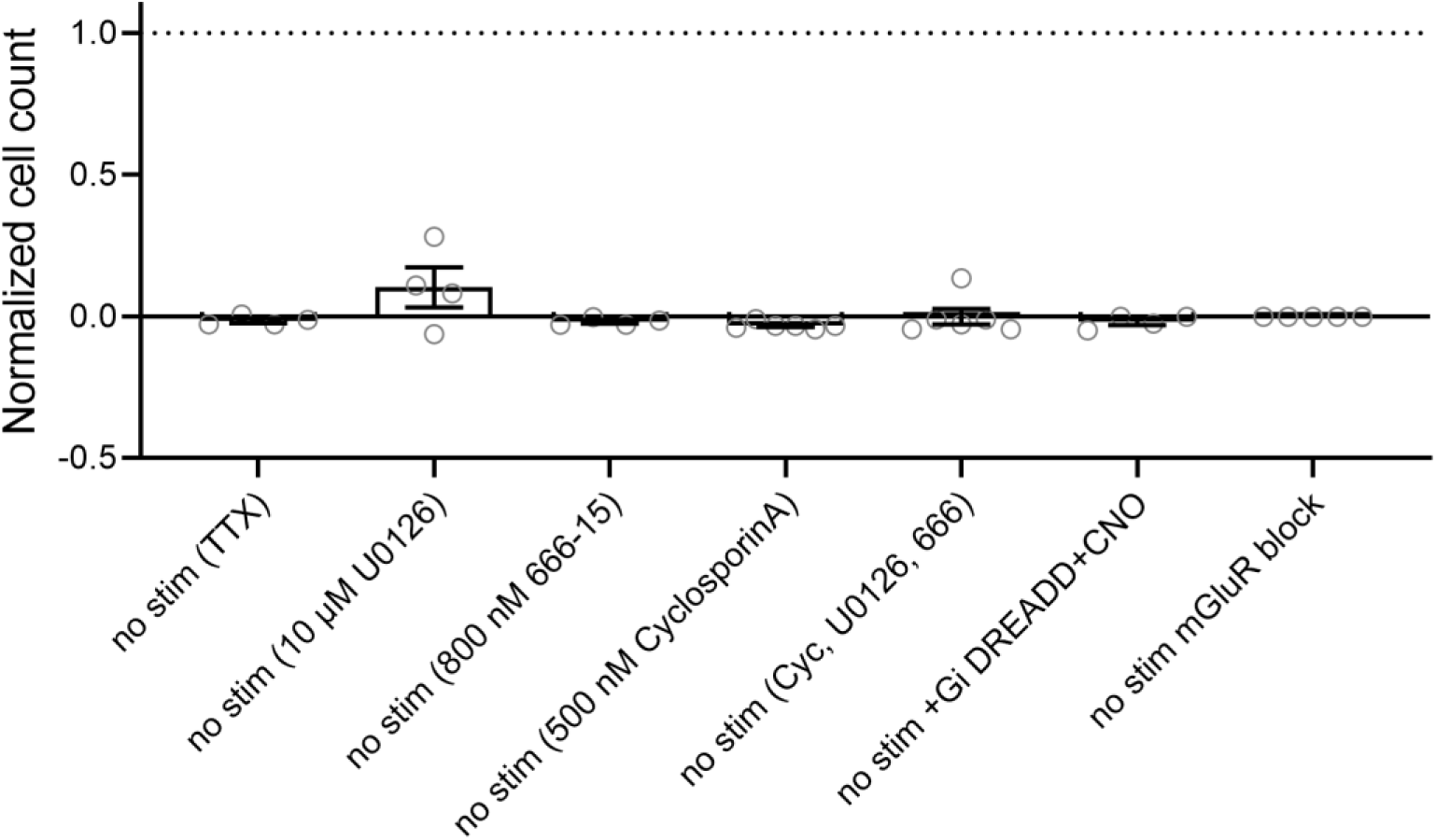
No effect of pharmacological agents tested on non-stimulated slice cultures. FOS+ neurons were counted in slice cultures (circles) incubated with blockers, counts were normalized to positive controls (high K^+^). Bars show mean ± SEM.

**Figure S5:**
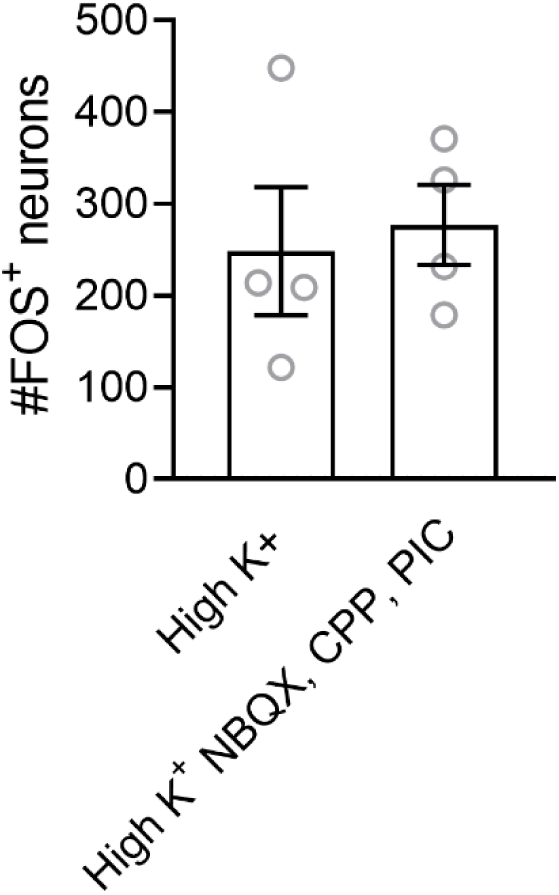
Inhibitors of fast synaptic transmission (NBQX, CPPene, picrotoxin) have no effect on high K^+^ induced FOS. High K^+^: n = 4 hippocampal slice cultures; High K^+^ with synaptic blockers: n = 4 hippocampal slice cultures. Bars show mean ± SEM.

